# Anthropogenic effects, not climatic, shaped Holocene population expansion of an insular bee fauna

**DOI:** 10.64898/2026.06.29.735417

**Authors:** Patricia S. Slattery, James B. Dorey, Bruno Alves Buzatto, Mark I. Stevens, Michael S.Y. Lee, Michael P. Schwarz

## Abstract

**Aim:** Remote island systems with small landmasses and reliable estimates of human occupancy are ideal model systems to disentangle the roles of global climatic changes and local human occupation on biota. Here, we tested this with demographic reconstructions of native, ground-nesting bees with non-specialised floral visitation habits to infer changes in effective population size (*N*_e_).

**Location:** Viti Levu, Fiji, southwest Pacific.

**Taxon:** *Lasioglossum* (*Homalictus*) bees (Hymenoptera: Halictidae)

**Methods:** We applied a combination of both mitochondrial and genomic datasets from five species distributed across the elevational range for our analyses. Both kinds of data were used for validation of population assignments and testing for structure, and to investigate any demographic shifts.

**Results:** All lowland species and populations showed strong signals of increasing *N*_e_ that correspond to the timing of human occupation of Fiji, but not Holocene climatic changes. Highland populations, with greater isolation and less affected by anthropogenic impacts, did not show evidence of recent rapid increases in *N*_e_. Population expansion rates across the elevational gradient differed between taxa, with earlier and larger increases in predominantly lowland species. This is consistent with the movement of people inland and into montane elevations of the island, and corresponding landscape changes that benefit the ecology of these bees.

**Main conclusions:** Specific life history traits of these bees, combined with substantive clearing of forest cover and floristic changes at lower elevations, has likely increased nesting opportunities and abundance of invasive floral resources. Our findings contrast with recent evidence that human occupation of Fiji has resulted in decreased ant biodiversity and raise the paradoxical possibility that human-mediated environmental changes may benefit some native montane tropical insect faunas.

## Introduction

There is an ever-increasing interest in predicting how biota will respond to changing environments (Alegría et al., 2022; Graham et al., 2017). One promising approach is examining how species and populations have responded historically, to infer how those responses might inform predictions for future climates (Nadeau et al., 2017; Wiens & Zelinka, 2024; Willis & MacDonald, 2011). Disentangling these population pressures is a persistent issue in evolutionary ecology, and major complexities arise when potential causal agents overlap in time, and discriminating between them can be complex (Field et al., 2008; van der Kaars et al., 2017; Wroe et al., 2013). Anthropogenic impact and climatic changes are well discussed drivers of demographic patterns, often with extensive chronological overlap (Archer et al., 2025; Saltré et al., 2016, 2019; Selwood et al., 2015). Demographic reconstructions for these populations can also rely heavily on the fossil record, which is problematic when it is comparatively incomplete and strongly biased for certain taxa (Schachat & Labandeira, 2021).

In instances where the fossil record is scarce, the application of molecular data to these questions opens alternative avenues to look backwards in time to explore selective pressures. Issues in delineating between drivers can be at least partially ameliorated in remote islands or other insular systems through limiting potential responses (Delgado et al., 2017; Russell & Kueffer, 2019). Patterns of immigration and emigration are limited by the dispersal capacity of the taxon, landmass area, and degree of isolation. In addition, there is often a much better understanding of human presence on islands, with archaeological information of initial settlements with increased detail following further colonisation (Rick et al., 2013; Russell & Kueffer, 2019).

At a population level, ectothermic species can respond to warming climates in insular systems by persisting within refugia, migrating to cooler climates, or adapting within their current range (González-Tokman et al., 2020; Lancaster et al., 2022; Sunday et al., 2014). Due to the constraints of insularity, warming climates and niche conservatism in more topographically complex systems push taxa to increasingly higher elevations to track favourable conditions (Chen et al., 2011; Linck et al., 2021; Maihoff et al., 2023; Marshall et al., 2020; McCain & Garfinkel, 2021). Alternatively, an adaptive response to change requires a degree of plasticity that can keep pace with environmental changes (Hoffmann et al., 2013; Hoffmann & Sgrò, 2011; Kellermann et al., 2012; Martin et al., 2023), which is unlikely for many ectotherms, especially those with less genetic load in the population (Kellermann & Heerwaarden, 2019; Matuszewski et al., 2015). Taxa facing these restrictions will be subject to ongoing pressures through climate-driven changes in community structure (Ponti & Sannolo, 2023; Shah et al., 2020), physiological function (Dahlhoff et al., 2019), genetic diversity, and effective population size (Arenas et al., 2012; Meza-Joya et al., 2023, 2025; Rogan et al., 2023).

Viti Levu is the largest island in the tropical archipelago of Fiji — an insular and vulnerable system with a relatively complete archaeological record. The earliest evidence of humans in Fiji comes from the Lapita people on Viti Levu, in the southwest of the island in the Bourewa region. Radiocarbon dating of marine shells and charcoal samples from human habitation are estimated to be ∼3,000 years old (Clark & Anderson, 2009b). Human populations were still very labile until the formation of more stable communities in the lowlands ∼2,500 years ago (Clark & Anderson, 2009a). The earliest evidence of movement into higher elevations is estimated at 2,100 years ago, but there is no evidence of fortified settlements until ∼1,300 years ago (Clark & Anderson, 2009a). Greater human presence and impacts in the lower elevations broadly fits with the forest cover and demographic patterns of Viti Levu today (Avtar et al., 2022; Fiji Bureau of Statistics, 2024). Substantial changes have occurred in the floristic makeup of Fiji since the arrival of the Lapita people (Hope et al., 2009), and the landscape of the archipelago is dominated by introduced plant taxa after European colonisation (Ash, 1992).

Fiji’s native *Lasioglossum* (*Homalictus*) bees provide an almost ideal model taxon to compare anthropogenic and climatic drivers of change in demography and geographic range. These Fijian bees are proposed as the product of a single dispersal event followed by major diversification (Groom et al., 2013), and consequent speciation largely driven by glacial, and interglacial climatic cycles (Dorey et al., 2020). The greatest diversity is present in higher elevations (Slattery et al., 2025), with phylogenetic and physiological limitations to dispersal outside of their climatic niche (da Silva et al., 2025; Dorey et al., 2020). As ground nesting bees, *Lasioglossum* in Fiji have a heavy reliance on cleared habitat (Ibalim et al., 2020) that *in situ* can be a result of natural land slips that increase in response to anthropogenic disturbance, as well as village-scale clearing for agriculture (Restrepo et al., 2009).

The intersection of climate and human presence on demography has been explored across the Fijian archipelago in the endemic bee *Lasioglossum fijiense*. The population expansion of this species fits well with records of human presence and consequent environmental modifications (Dorey et al., 2021). However, that study was species-specific, and focussed on a supergeneralist (Draper et al., 2021) bee with a mostly low elevational range, and may therefore not effectively represent congenerics with different elevational niches (da Silva et al., 2025; Slattery et al., 2025), or the changes in land use across this elevational gradient (Avtar et al., 2022).

Ectothermic taxa have well described responses to a warming climate, often dependent on their niche use along the elevational gradient (García-Robledo et al., 2016; Shah et al., 2020; Warren et al., 2018). Warmer-adapted low elevation species often experience population growth as their thermal niche expands up the mountain, while cool-adapted higher elevation species with more specialised thermal niches would experience range contractions with population declines as their thermal niche is pushed to increasing elevations (da Silva et al., 2025; McCain & Garfinkel, 2021). We hypothesised similar patterns to be retrieved in the demography of other Fijian *Lasioglossum* species if climatic variables were a major driver. Alternatively, if the population trends of these bees were not primarily driven by climate, population expansion would be more closely linked to human presence (Dorey et al., 2021).

To test the applications of insular systems in disentangling these divers, we aimed to infer historical demographic patterns of *Lasioglossum* (*Homalictus*) *fijiense*, *L. tuiwawae*, *L. groomi*, *L. ostridorsum* and an undescribed species from Dorey *et al*. (2020), ‘*L.* sp. *S*’, from across the elevational gradient of Viti Levu. Mitochondrial DNA from the cytochrome *c* oxidase subunit 1 gene (COI), supplemented by genome-wide single nucleotide polymorphisms (SNPs), were used to assess the contemporary structure and historical demography of populations of native *Lasioglossum*. We discuss our results with the aim of exploring if demographic changes in these bees correspond most closely to historical climatic changes or the impacts of human occupation.

## Materials and Methods

### Specimen collection and preparation

Across multiple seasons from 2010 to 2018, researchers collected five species of *Lasioglossum* via sweep netting from the main island of Fiji, Viti Levu. We collected specimens of *Lasioglossum* (*Homalictus*) *fijiense*, *L. groomi*, *L. ostridorsum*, *L. tuiwawae* and *L.* sp. *S* across the climatic and elevational range of Viti Levu (0–1,328 m ASL; above sea level), with the density of collections varying geographically, based on accessibility (Figure 1). Each collection recorded GPS coordinates and elevation, with specimens preserved in >98% ethanol — which was replaced within one week of collection — and stored at ∼ 5°C for DNA preservation.

**Figure 1:**
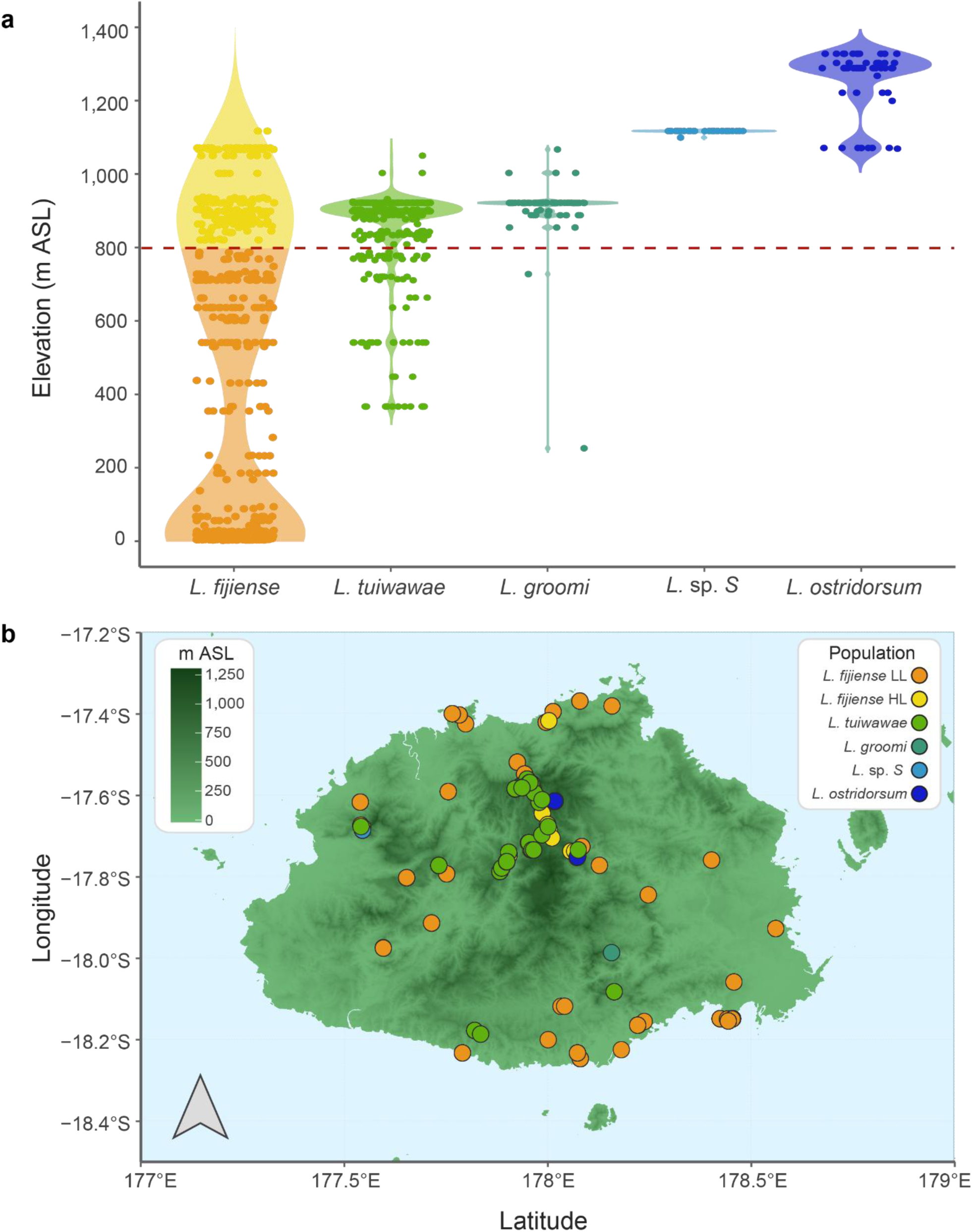
**a)** Violin plot showing the collection density of COI-sequenced specimens of five Fijian *Lasioglossum (Homalictus)* species distributed across the elevational gradient on Viti Levu. Width of the violin indicates a greater density of samples, with the red dashed line distinguishing between highland and lowland elevations. **b)** Viti Levu’s digital elevation model (sourced via https://srtm.csi.cgiar.org/; Reuter et al., 2007) was overlaid with specimen collection locations. Point colours correspond to population assignments, with increasingly darker green of the island corresponding to increased elevation. *L. fijiense* lowland (LL) population individuals are represented by orange circles, and highland (HL) individuals are represented by yellow circles.

Tissue samples for mitochondrial (mt) DNA cytochrome *c* oxidase subunit 1 (COI) extraction and amplification were obtained by the removal of one hind leg for sequencing. Samples collected prior to 2015 were sequenced at the Canadian Centre for DNA

Barcoding at the Biodiversity Institute of Ontario with primers LepF1 and LepR2 (Hebert et al., 2004). From 2015 onwards, *COI* was extracted at the South Australian Regional Facility for Molecular Ecology and Evolution using a Gentra Puregene® DNA Purification kit, with sequencing performed at Macrogen, Inc. (Korea) with primers LCO1490/ HCO2198 or LepF1/LepR2.

For genome-wide single nucleotide polymorphisms (SNPs), DNA was extracted from the thorax (mesosoma), fore- and midlegs of only female specimens — to avoid any confounding effects of male haploidy on SNP genotypes. Tissue samples were sent to Diversity Arrays® (Australia) for SNP assays through high throughput microarray sequencing (DArTseq) via the restriction enzymes PstI and Hpall (Jaccoud et al., 2001). Single nucleotide polymorphisms were identified based on pooled-species samples in initial analyses, so many SNP alleles may be unique to only one or a few species.

### Alignment creation and SNP filtering

*COI* sequences were compared against the NCBI BLAST database for non *Lasioglossum* sequences and aligned in Geneious Prime 2020.1.1 (http://www.geneious.com/). The *COI* sequences were aligned and placed into the correct reading frame and primers and poor sequence trimmed for all species through visual inspection, resulting in a final alignment of up to 630 base pairs. We used R (version 4.3.3; R Core Team, 2024) and the RStudio environment (version 2024.04.1+748; Posit Team, 2024) to calculate pairwise *F*_ST_ values based on Nei (1989) with the package *hierfstat* (version 0.5-11; Goudet, 2004). Elevational population structure was detected in *L. fijiense* (Supplementary Table 1 in Supporting Information), and as such, the species was treated as two populations in subsequent *COI* analyses to avoid violating assumptions of extended Bayesian skyline plots (eBSP) and mismatch analyses (Grant, 2015; Heller et al., 2013). Populations of *L. fijiense* were split dependent on collection elevation, with highland specimens occurring > 800 m ASL, and lowland specimens below this boundary. This demarcation has been described, and coincides with the elevation where cloud forests and associated floristic change begins (Ash, 1992; Dorey et al., 2020; Staines et al., 2017). Final *COI* population alignments were as follows: lowland *L. fijiense* (n = 369; 627 bp), highland *L. fijiense* (n = 204; 627 bp), *L. tuiwawae* (n = 452; 630 bp), *L. groomi* (n = 67; 630 bp), *L.* sp. *S* (n = 36; 630 bp) and *L. ostridorsum* (n = 50; 630 bp).

SNPs were filtered in R via the *dartR* package (version 2.9.7; Mijangos et al., 2022) from 62,426 SNPs sequenced across five species (n = 94 total). Species were independently filtered for locus call rates with a threshold of 0.95, reproducibility with a threshold of 0.99, monomorphs, secondaries and any missing loci. After filtering, *L. fijiense* (n = 19) had 3,768 SNPs, *L. tuiwawae* (n = 19) had 4,611 SNPs, *L. groomi* (n = 19) had 3,018 SNPs, *L.* sp. *S* (n = 19) had 1,566 SNPs, and *L. ostridorsum* (n = 18) had 2,340 SNPs. Due to the similarity in genomic signature (Supp. Fig. 2) and small number of individuals that represent *L. fijiense*, populations were pooled for genomic analyses.

**Figure 2:**
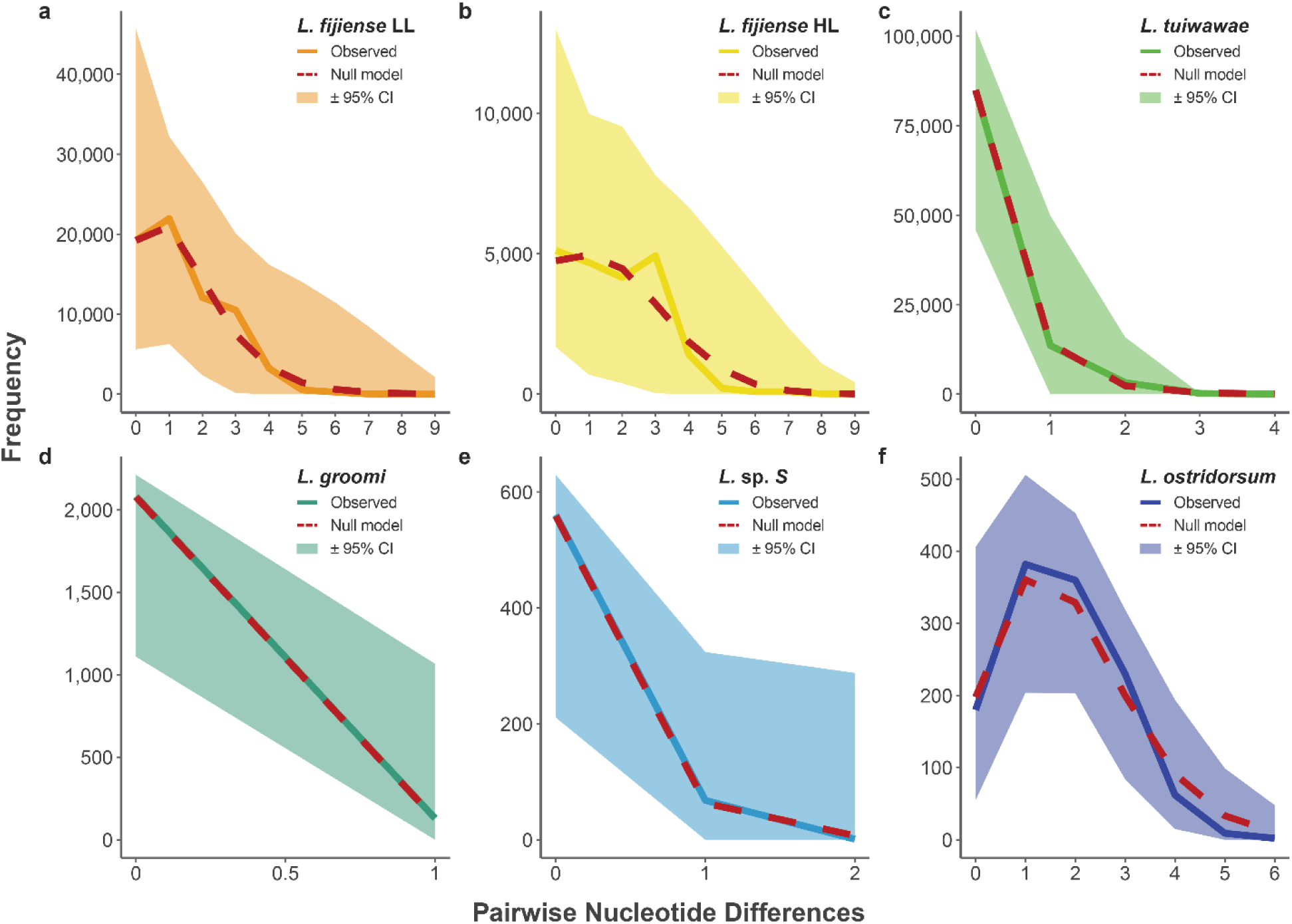
Mismatch analyses for populations of Fijian *Lasioglossum (Homalictus)*, calculated using COI data under a null model of recent population expansion. Mismatch frequency is presented along the *y* axis, and nucleotide differences on the *x* axis, while the simulated null model for each population is denoted by the red dashed line, and the solid-coloured line is observed differences with the shaded area representing 95% confidence intervals of the observed values. **a)** *L. fijiense* lowland (LL) population is denoted by the solid orange line (n = 369; 627 bp), **b)** solid yellow represents the highland (HL) population of *L. fijiense* (n = 204; 627 bp), **c)** the green line for *L. tuiwawae* (n = 452; 630 bp), **d)** the teal line representing *L. groomi* (n = 67; 630 bp), **e)** the light blue denoting *L.* species S (n = 36; 630 bp) and **f)** the dark blue representing *L. ostridorsum* (n = 50; 630 bp).

### Population metrics

We used mismatch analyses in Arlequin (version 3.1; Excoffier et al., 2007) to compare the observed and predicted haplotype frequencies in *COI* across *Lasioglossum* populations. The analyses ran for 16,000 simulations under a null hypothesis of recent population expansion and the outputs were then imported into R and plotted with *ggplot2* (version 3.5.2; Wickham, 2016). Arlequin was also used for inferring sample characteristics and diversity, along with exploring variance within and between populations (described in Supp. Tables 3 and 4). Any populations with insufficient specimens or haplotypic diversity (*i.e.*, less than four nucleotide differences and/or less than 50 specimens) were excluded from eBSP analyses, due to inadequate inferential power. PopART (version 1.7; Leigh & Bryant, 2015) minimum-spanning networks were created to visualise the relationship between populations.

Genomic SNP populations were assessed with *dartR* in R, using Gower principal coordinates analysis (PCoA) to assess and validate species assignments, with Bonferroni-corrected ternary plots visually inspected for significant deviations from Hardy-Weinberg equilibrium (Supp. Fig. 7). To confirm that changes in effective population size based on *COI* data were not driven by mitochondrial sweeps, we tested genomic SNP data for deviations of neutrality in mutations with Tajima’s *D* statistic (Tajima, 1989) using the package *pegas* (version 1.3; Paradis, 2010).

### Extended Bayesian skyline plots

Historical demographic analyses were run in BEAST2 (version 2.6.7; Bouckaert et al., 2019) under a coalescent eBSP tree prior (Heled & Drummond, 2008). Those mtDNA populations that had sufficient genetic diversity and representative specimens for the analyses were the *L. fijiense* highland and lowland populations, *L. tuiwawae, L. groomi* and *L. ostridorsum*. Analyses of *COI* were restricted to the 3^rd^ codon position (non-coding in Hymenoptera) to avoid violating the assumption of neutrality in genetic markers. These restricted alignments were run across an optimised length, number and temperature of chains via a Metropolis-coupled Markov chain Monte Carlo algorithm with the package *CoupledMCMC* (Müller & Bouckaert, 2020; see Supp. Table 5 for specifics and executables). All *COI* populations used a HKY substitution model (Hasegawa et al., 1985) under a strict clock, with a population factor of 0.5 (since only female mtDNA contributes to effective population size); our data were insufficient for more complex models and would not achieve convergence.

Log files from the eBSP analyses were visually inspected for convergence in Tracer (version 1.7; Rambaut et al., 2018) and run through the *EBSPAnalyser* tool provided in the BEAST2 applications. This output was modelled in R with *ggplot2*, and axes converted to chronological time and recognisable units in Adobe Illustrator (version 29.5.1; Adobe, 2025). Mean sea surface temperature was calculated from Stott *et al*. (2004) using records from four sites in the western Pacific Ocean, and plotted with *ggplot2*.

The axes in eBSPs were adjusted based on the estimated mutation rate of *Drosophila melanogaster* with a high A/T bias across the mitogenome (82%; Haag-Liautard et al., 2008), and *Caenorhabiditis elegans* with a more similar A/T bias in the mitochondrial third codon position (70.3%) to that of *L. fijiense* (Denver et al., 2000). As BEAST2 ignores indels in estimation, we corrected for their removal in the rate presented by Haag-Liautard *et al*. (2008) with a final adjusted mutation estimate of 4.8 x 10^8^ per generation. Including both allows a more nuanced approach with a more comparable taxon, while still appropriate for direct comparison to previously published results in related studies (Dorey et al., 2021). Axes were converted to years before present under the assumption of four generations per year (Dorey et al., 2021; Groom et al., 2014).

## Results

### Population metrics

We found evidence of population structure across elevation for *L. fijiense* (*F*_ST_ = 0.12; Supp. Table 1), with both populations showing limited gene flow and haplotypes (Supp Table 3 and Supp. Fig. 6). This result, combined with our analysis of molecular variance (AMOVA; Supp. Table 4), validated our population assignments. The greatest haplotypic diversity of any population was present in *L. ostridorsum* (H = 0.85 ± 0.02; Supp Table 3 and Supp. Fig. 6), and the least present in *L. groomi* (H = 0.06 ± 0.04; Supp Table 3 and Supp. Fig. 6). We used mismatch analyses and minimum-spanning haplotype networks to test assumptions of recent population expansion. All *COI* populations showed a good fit to the null model of recent population expansion in mismatch analyses (Figure 2), with the highland population of *L. fijiense* and *L. ostridorsum* suggesting the strongest deviations, though not significantly so.

**Table 1:** Tajima’s *D* statistics used to determine deviations from neutral mutations in five species of Fijian *Lasioglossum (Homalictus)* bees using SNP data. Calculated using the package *pegas* (Paradis, 2010) from methods described in Tajima (1989) with an α value set to 0.05. Significant deviations from expected polymorphisms under equilibrium are indicated with an asterisk (*).

| | Tajima's *D* | p-value (normal dist.) | p-value ($\beta$ dist.) |
| --- | --- | --- | --- |
| *L. fijiense* | -2.135 | 0.03 * | 0.01 * |
| *L. tuiwawae* | -2.208 | 0.03 * | 0.01 * |
| *L. groomi* | -1.927 | 0.05 * | 0.03 * |
| *L. sp. S* | -2.037 | 0.04 * | 0.02 * |
| *L. ostridorsum* | -1.877 | 0.06 | 0.04 * |

Our genomic data suggested a similar trend of increasing population, with negative Tajima’s *D* statistics for all populations (Table 1). Of all species, *L. ostridorsum* was the only that didn’t significantly deviate from equilibrium under both distributions (Table 1). A PCoA of filtered SNP loci supported five clear species groupings (Supp. Fig, 2), and Bonferroni-corrected ternary plots suggested no significant deviations from Hardy-Weinberg equilibrium outside of *L.* sp. *S* (Supp. Fig. 7).

### Extended Bayesian skyline plots

In all cases, our effective population (*N*_e_) estimates of *Lasioglossum* did not show any fluctuations consistent with climatic variables under either of the proposed mutation rates in the eBSPs (Figure 3). The populations of lowland *L. fijiense*, *L. tuiwawae* and *L. groomi* (Figure 3) showed stronger increases in *N*_e_ closer to the present. Highland *L. fijiense* showed little evidence of any demographic change (Figure 3), and *L. ostridorsum*, at the highest elevation, had a stable increase in *N*_e_ (Figure 3) as far back as we can reliably infer. Median declines closest to the present are likely a result of bias or structure and small relative to the *y*-axis while corresponding to increases in 95% highest posterior densities, so should not be interpreted (Heller et al., 2013).

**Figure 3:**
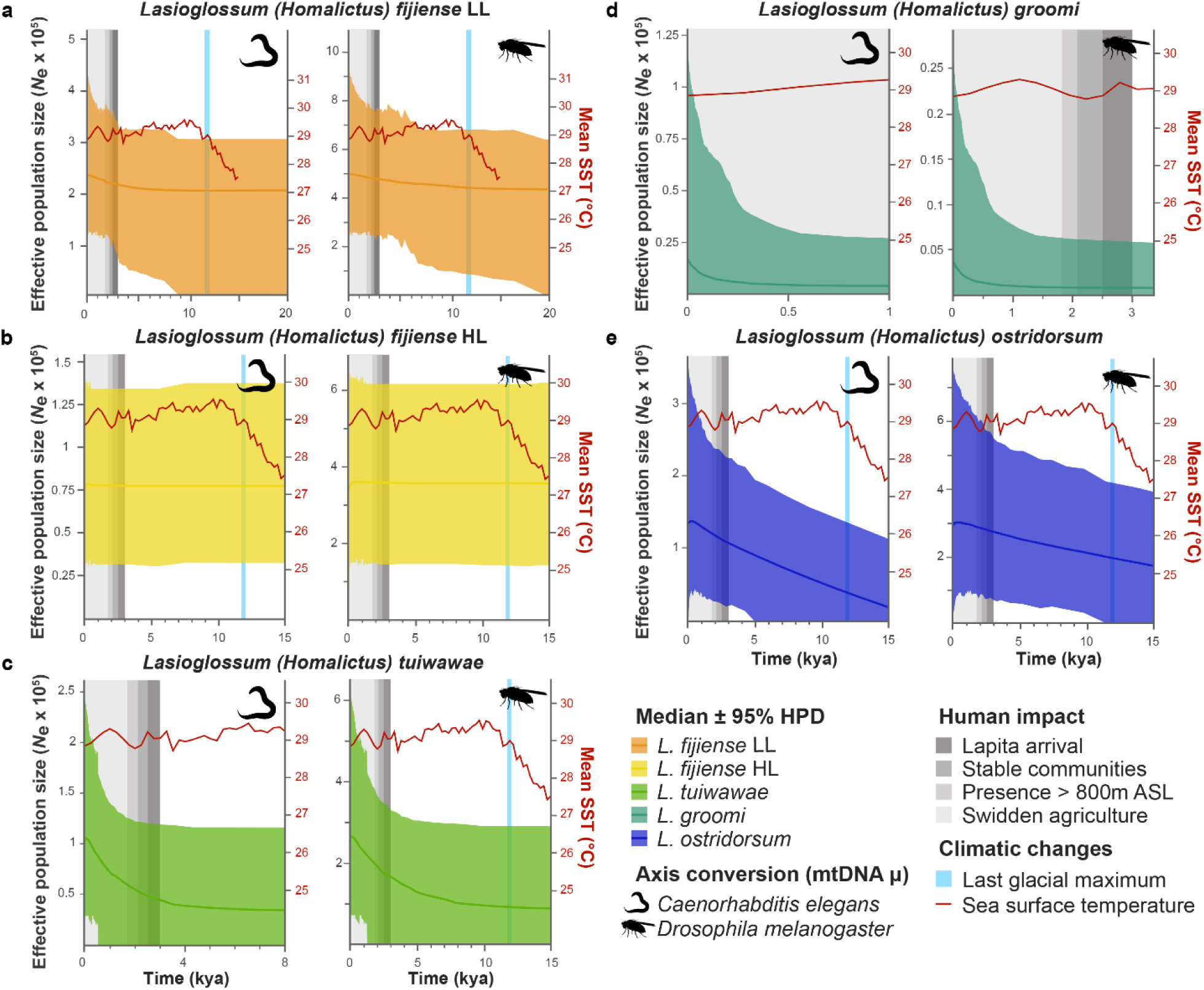
Extended Bayesian skyline plots for five populations of Fijian *Lasioglossum (Homalictus*), estimated based on 3^rd^ codon positions of mtDNA COI. Each panel shows historical demographic patterns of a population under two alternative mutation rates (Denver et al., 2000; Haag-Liautard et al., 2008), used to convert the *x* and *y* axes, which detail time since the present and effective population size (*N*_e_), respectively. Dates of interest relating to human impact are denoted in shades of grey, with the average sea surface temperature overlaid in red (based on Stott et al., 2004), and the end of the last glacial maximum in light blue. Median demographic trends are an opaque line, with a semi-opaque 95% highest posterior density (HPD) coloured **a)** orange for lowland *L. fijiense* (n = 369; 627 bp), **b)** yellow for highland *L. fijiense* (n = 204; 627 bp), **c)** green for *L. tuiwawae* (n = 452; 630 bp), **d)** teal for *L. groomi* (n = 67; 630 bp), and **e)** dark blue for *L. ostridorsum* (n = 50; 630 bp).

## Discussion

We used both mitochondrial and genomic data across multiple analyses to explore demographic trends in Fijian *Lasioglossum* (*Homalictus*) bees on Viti Levu. Distributed across the entire elevational gradient, our taxa likely vary in their use of climatic niche space and exposure to human impacts, but all analyses supported effective population increases in every group. In a partial rejection of our hypothesis, there was no evidence of demographic fluctuations or declines in highland populations that aligned to either of our climatic conditions of interest (*i.e.* mean sea surface temperature or last glacial maximum). Instead, patterns of population expansion in *Lasioglossum* matched better with archaeological evidence of human presence, and changes in anthropogenic impact with increasing elevation.

Recent research in Fiji suggests divergent responses in demography under anthropogenic modification, with the diversity of endemic ants in decline (Liu et al., 2025), contrasting to population increases of *L. fijiense* across the archipelago (Dorey et al., 2021). Dorey et al. (2021) focussed on demographic reconstructions of a primarily lowland bee, while most endemic ants in Liu et al. (2025) are restricted to higher elevations. To address this conflict, we extended the research by Dorey et al. (2021) by including a greater number of endemic species from a broader elevational range, and more nuance through the application of multiple mutation rates and validated population assignments.

We used both mitochondrial and genomic data to test population groupings and species assignments by estimating population structure and differentiation. Consistently, we found support for five species of *Lasioglossum*, though with significant population structure in mtDNA for highland and lowland *L. fijiense* (see Supporting Information). Results of pairwise mismatch analyses and extended Bayesian skyline plots (eBSP) provide evidence of population expansion in all species from mitochondrial data (Figures 2 and 3). Additionally, our Tajima’s *D* values using genomic SNPs provided further evidence that these patterns reflect true population expansion, rather than the aftermath of a mitochondrial sweep/bottleneck (Table 1).

When looking at populations individually, the timing and degree of this increase differs along the elevational gradient, along with intensity and timing of human presence and associated environmental modification (Figure 3). The arrival of the Lapita people is the earliest evidence of human presence on the island (∼ 3,000 years ago; Clark & Anderson, 2009b), but evidence of stable communities, movement into the highlands and agricultural intensification occurred in the thousand or more years following (Clark & Anderson, 2009a; Roos et al., 2016). Currently, the highest Fijian elevations still host comparatively few permanent human settlements, and much greater forest cover (Avtar et al., 2022; Fiji Bureau of Statistics, 2024).

Demographic reconstructions for lowland *L. fijiense* produced effective populations that coincided with the earliest evidence of humans on Viti Levu, while steeper median curves in *L. tuiwawae* aligned with more recent human impact (Figure 3). Notably, *L. ostridorsum*, at the highest elevation with the least exposure to human impact, and had the greatest haplotypic diversity, and showing a smooth, continuous demographic growth. None of the highland populations in this study showed any evidence of the population declines we might expect under a range contraction driven by climate (Dahlhoff et al., 2019; Shah et al., 2020), nor the negative demographic consequences of anthropogenic pressures (Liu et al., 2025; Russell & Kueffer, 2019), with trends suggesting stable or closer to exponential increases in effective population. This positive trend occurs across the elevational gradient, matching historical human occupancy patterns, and more recent infrastructural changes (Avtar et al., 2022; Clark & Anderson, 2009b).

The widespread habitat available to *L. fijiense* in the lowlands is primarily driven by human modification for agricultural practices and/or the development of related infrastructure (Davies et al., 2024). Fiji has a multitude of invasive flora present across the island, much of which was introduced for ornamental, forestry or agricultural purposes (Lowry et al., 2020; Meyer, 2014). Native supergeneralist pollinators are well documented in islands systems (Olesen et al., 2002; Picanço et al., 2017), and include *L. fijiense* in Fiji (Draper et al., 2021; Hayes et al., 2019; Staines et al., 2017). Using a wide variety of both introduced and native floral resources, *L. fijiense* has a preference for nesting in open areas, and would likely benefit from these anthropogenic changes to the environment (Draper et al., 2021; Hayes et al., 2019; Ibalim et al., 2020). Previously inferred increases in the effective population size of *L. fijiense* was therefore attributed to anthropogenic modifications (Dorey et al., 2021), and shared life-history traits of related species could predict similar responses to these human impacts.

We found that the demographic trends of these Fijian bees fit better with records of human occupation, rather than Holocene patterns in climate change. Further studies in other tropical and insular pollinator suites would reveal if this is a broad pattern. If such patterns are indeed widespread, it raises the question of how ‘natural’ plant-pollinator ecosystems are, even before European colonisation. Based on the foundational goals of conservation: should strategies attempt to achieve preservation and/or rehabilitation of pre-colonial or pre-human ecosystems? In turn, if these prior structures are already disrupted in some manner (either positively or negatively), to what extent is an insular ecosystem regarded as ‘natural’?

## Data Accessibility Statement

All genetic and genomic datasets with associated metrics files are available on Figshare, along with R scripts and exemplar XML code (https://figshare.com/s/a36f5e2ea8fbf85702a6).

## Notes

https://doi.org/10.25451/flinders.31417679

### Competing Interest Statement

The authors have declared no competing interest.

### Funding

- Australian Federal Government New Colombo Plan: NCPST Fiji 15482

