## Supplementary figures and tables for "Anthropogenic effects, not climatic, shaped Holocene population expansion of an insular bee fauna"

**Supplementary Table 1:** Pairwise $F_{ST}$ for mtDNA COI data (630 bp) from six populations of Fijian *Lasioglossum (Homalictus)* bees. Calculated using Tajima & Nei's method for distance matrix creation. *L. fijiense* populations are split into lowland (LL) and highland (HL). Asterisks ('*') indicate a significant p-value of < 0.05, based on 16,000 permutations.

| | | ***L. fijiense*** | | ***L. tuiwawae*** | ***L. groomi*** | ***L. sp. S*** | ***L. ostridorsum*** |
|---|---|---|---|---|---|---|---|
| | | LL | HL | | | | |
| ***L. fijiense*** | LL | | | | | | |
| | HL | 0.116* | | | | | |
| ***L. tuiwawae*** | | 0.973* | 0.976* | | | | |
| ***L. groomi*** | | 0.95* | 0.944* | 0.994* | | | |
| ***L. sp. S*** | | 0.966* | 0.961* | 0.991* | 0.998* | | |
| ***L. ostridorsum*** | | 0.959* | 0.952* | 0.985* | 0.979* | 0.936* | |

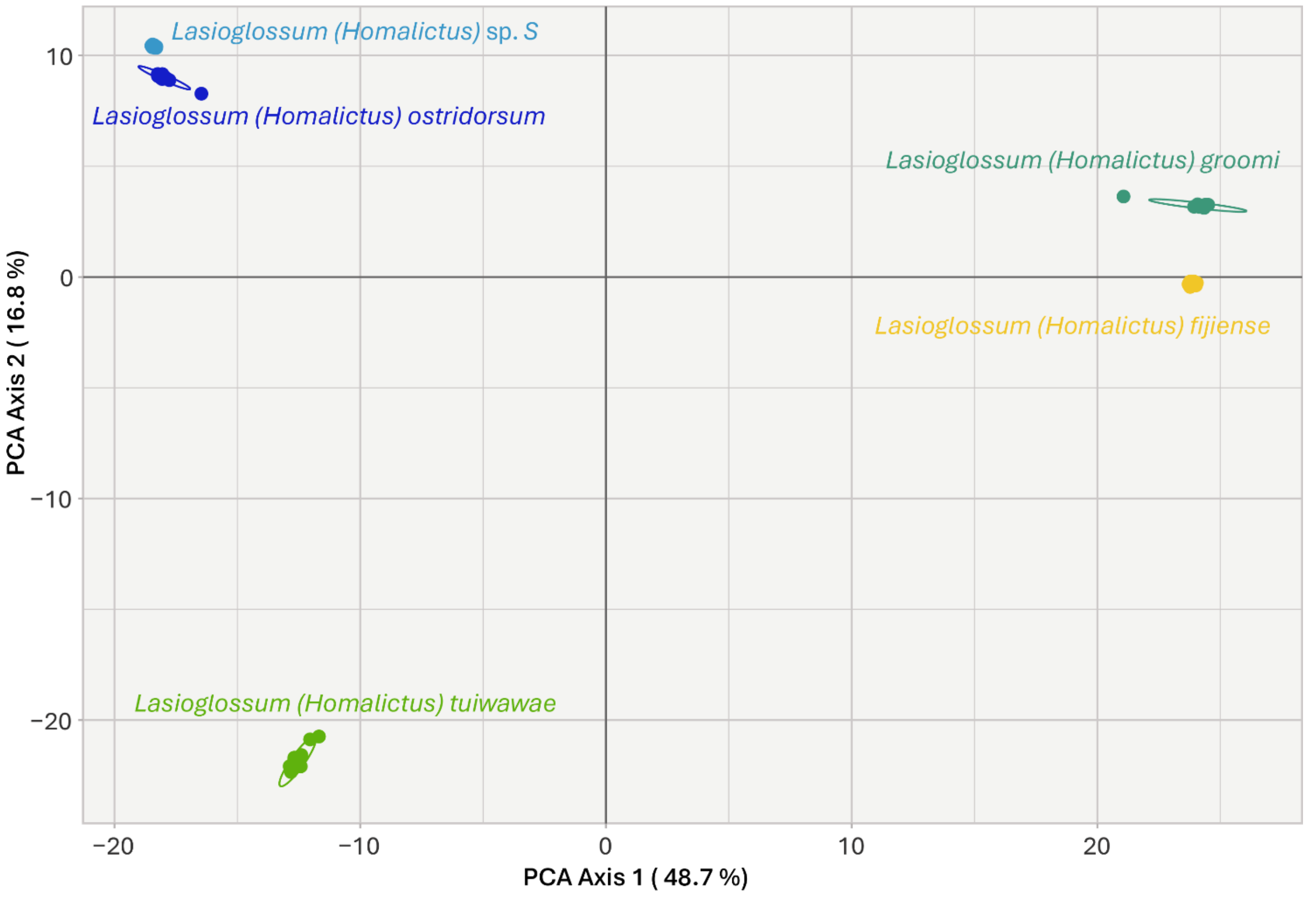


**Supplementary Figure 2:** Gower Principal Coordinates Analysis (PCoA) for five species of *Lasioglossum (Homalictus)* found on Viti Levu, Fiji. Yellow points are *L. fijiense* (n = 19), green for *L. tuiwawae* (n = 19), *L. groomi* (n = 19) points are teal, *L.* sp. *S* (n = 19) is light blue, and *L. ostridorsum* (n = 18) are dark blue. The PCoA is based on a combined filtered dataset of 8,381 SNPs from all 94 individuals.

**Supplementary Table 3:** Sample characteristics and molecular diversity summary of mitochondrial COI (630 bp) for six populations of *Lasioglossum* (*Homalictus*) bees. H = haplotype diversity, π = nucleotide diversity, θ = theta (*S*) are presented for each population (± 1 standard deviation). *L. fijiense* is split into lowland (LL) and highland (HL) populations due to population structure (see Supplementary 1)**.** Shared haplotypes of *L. fijiense* (n = 8) are listed, with clarification on equivalent specimen representation for each population.

| | | **No. ind.** | **No. haplo-types** | **No. poly-morphic loci** | **H (± SD)** | **Π (± SD)** | **Θ (*S*) (± SD)** | **Shared haplotypes (LL = HL)** |
|---|---|---|---|---|---|---|---|---|
| ***L. fijiense*** | LL | 369 | 32 | 29 | 0.716 (± 0.019) | 0.002 (± 0.002) | 4.471 (± 1.189) | F001 = F008,<br>F039 = F032,<br>F005 = F019,<br>F012 = F048,<br>F339 = F121,<br>F387 = F024,<br>F390 = F410,<br>F081 = F135 |
| | HL | 204 | 18 | 23 | 0.753 (± 0.016) | 0.003 (± 0.002) | 3.903 (± 1.149) | |
| ***L. tuiwawae*** | | 452 | 17 | 18 | 0.168 (± 0.024) | 0.0 (± 0.0) | 2.691 (± 0.803) | - |
| ***L. groomi*** | | 67 | 2 | 1 | 0.059 (± 0.039) | 0.0 (± 0.0) | 0.209 (± 0.209) | - |
| ***L. sp. S*** | | 36 | 3 | 2 | 0.11 (± 0.07) | 0.0 (± 0.0) | 0.482 (± 355) | - |
| ***L. ostridorsum*** | | 50 | 13 | 12 | 0.853 (± 0.026) | 0.003 (± 0.002) | 2.68 (± 1.045) | - |

**Supplementary Table 4:** Analysis of molecular variance (AMOVA) for within and between six populations of *Lasioglossum* (*Homalictus*) based on mtDNA COI (630 bp). Significance was based on 16,000 permutations with an alpha of < 0.05. *L. fijiense* is split into lowland (LL) and highland (HL) populations due to population structure (see Supplementary 1)**.**

| **Source of variation** | **df** | **Sum of squares** | **Variance components** | **Percentage of variation (%)** | ***p*-value** |
|---|---|---|---|---|---|
| Between populations | 5 | 222.57 | 0.262 Va | 53.09 | <0.001 |
| Within populations | 1,172 | 270.85 | 0.231 Vb | 46.91 | - |
| Total | 1,177 | 493.41 | 0.493 | - | - |

**Supplementary Table 5:** Extended Bayesian skyline plot parameters for each population of Fijian *Lasioglossum (Homalictus)* using third codon positions of mtDNA COI. *L. fijiense* mtDNA data is split into lowland (LL) and highland (HL) populations due to population structure (see Supplementary 1)**.**

<table>
<tr><th colspan="3">Population</th><th>Chain length</th><th>Chain number</th><th>Chain heat</th><th>Log rate</th></tr>
<tr><td rowspan="5">mtDNA COI data</td><td rowspan="2">L. fijiense</td><td>LL</td><td>1 billion</td><td>4</td><td>0.2</td><td>500,000</td></tr>
<tr><td>HL</td><td>500 million</td><td>8</td><td>0.2</td><td>500,000</td></tr>
<tr><td colspan="2">L. tuiwawae</td><td>500 million</td><td>8</td><td>0.1</td><td>500,000</td></tr>
<tr><td colspan="2">L. groomi</td><td>1 billion</td><td>4</td><td>0.2</td><td>500,000</td></tr>
<tr><td colspan="2">L. ostridorsum</td><td>1 billion</td><td>16</td><td>0.25</td><td>500,000</td></tr>
</table>

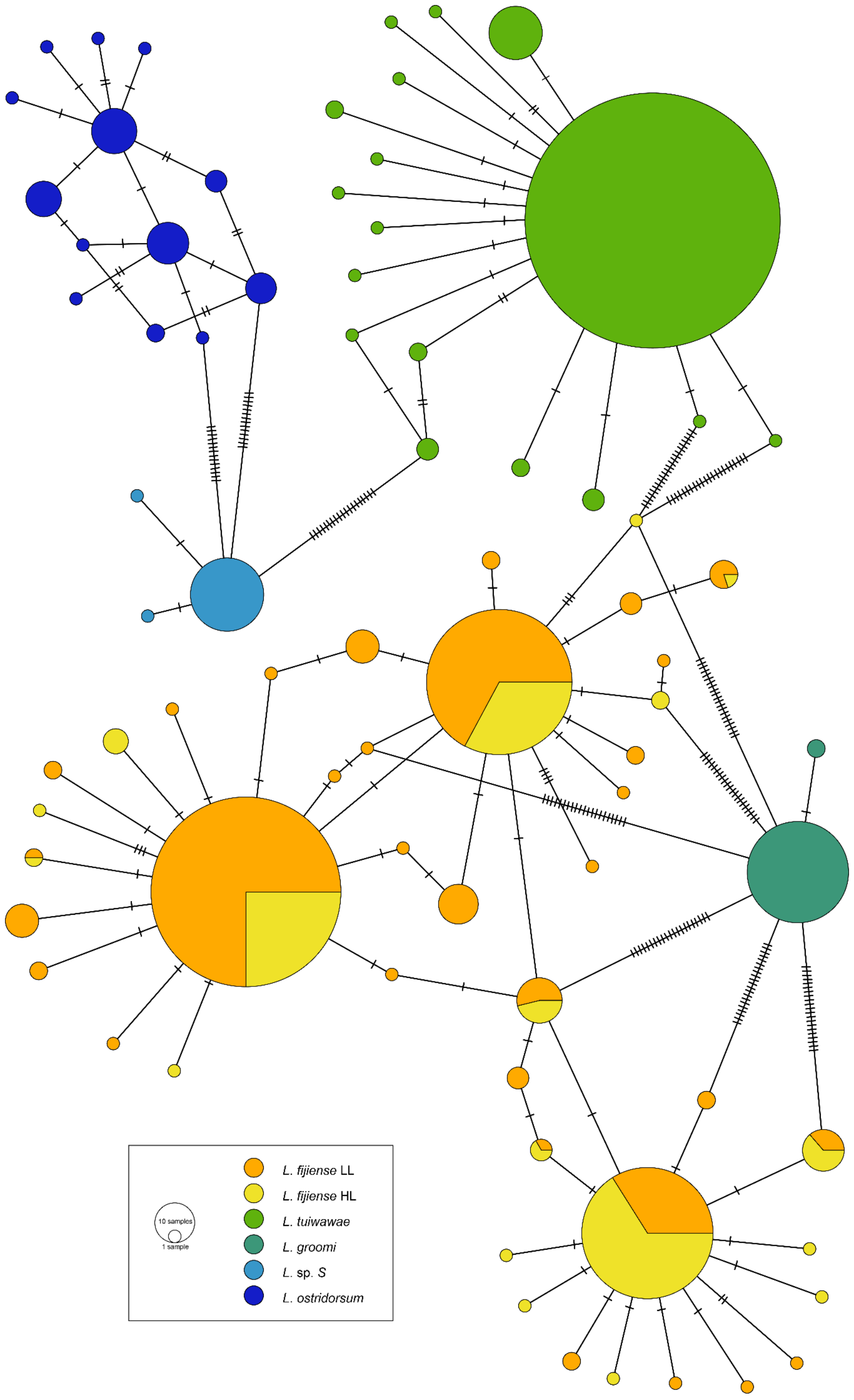
10 samples
1 sample
L. fijiense LL
L. fijiense HL
L. tuiwawae
L. groomi
L. sp. S
L. ostridorsum

**Supplementary Figure 6:** Minimum-spanning haplotype networks based on mitochondrial COI for five populations of Fijian *Lasioglossum (Homalictus)*. *L. fijiense* is split into lowland (LL) and highland (HL) populations due to population structure (see Supplementary 1). Size of the circles represents the number of sequences specimens with that haplotype (627 – 630 bp), with colour denoting which population the haplotype belongs to (Supplementary 2 summarises haplotype statistics for each population). Crosshatches between haplotypes indicate the number of nucleotide changes between each unique sequence.

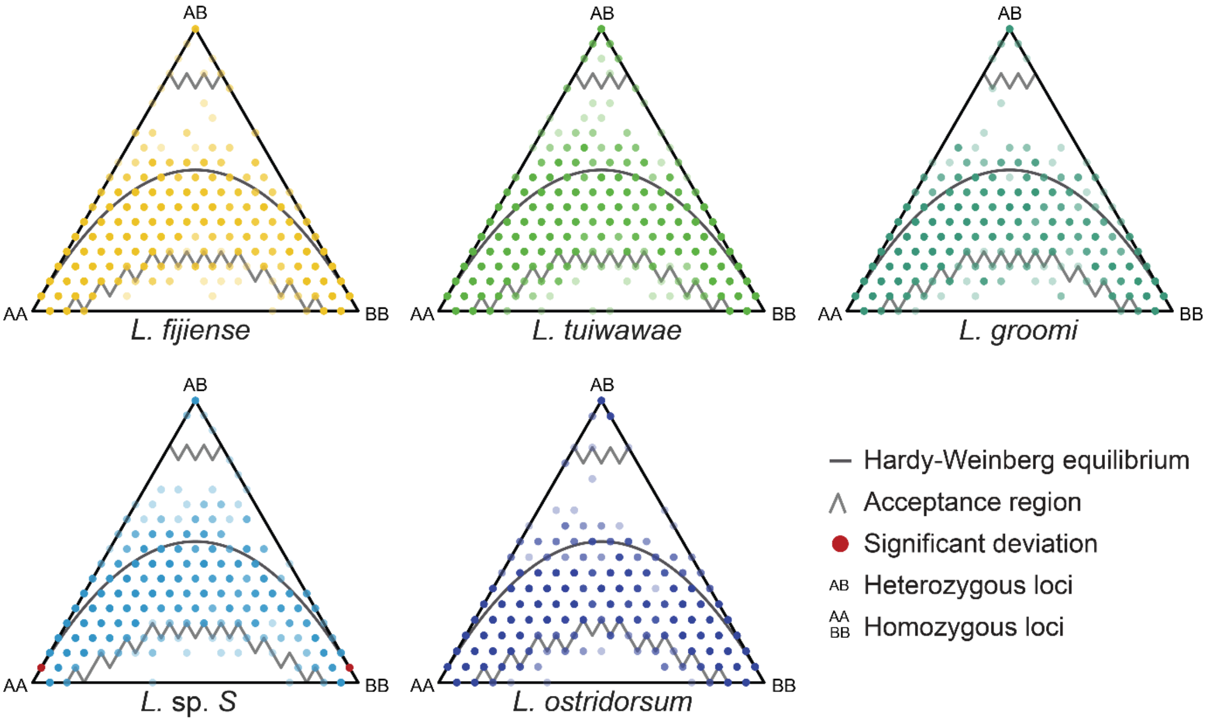


**Supplementary Figure 7:** Bonferroni-corrected ternary plots testing fit to Hardy-Weinberg equilibrium (HWE) for each species of Fijian *Lasioglossum (Homalictus)*. Both corners on the base of the triangle represent alternative homozygous loci, while the peak represents heterozygous SNPs. Testing for significant deviations to HWE used a Chi-squared test with an alpha value set to 0.05. The *L. fijiense* was calculated from 3,768 loci (n = 19), *L. tuiwawae* from 4,611 SNPs (n = 19), *L. groomi* from 3,018 loci (n = 19), 1,566 SNPs (n=19) from *L.* sp. *S* and 2,340 loci (n = 18) across *L. ostridorsum*. The darker grey parabolic curve represents the Hardy-Weinberg equilibrium with the accepted range denoted by lighter grey jagged lines and loci that have significantly deviated from this coloured red.
